# Comprehensive assessment of functional effects of commonly used sweeteners on *ex vivo* human gut microbiome

**DOI:** 10.1101/2022.01.15.474735

**Authors:** Zhongzhi Sun, Wenju Wang, Leyuan Li, Xu Zhang, Zhibin Ning, Janice Mayne, Krystal Walker, Alain Stintzi, Daniel Figeys

## Abstract

The gut microbiome composition and function are associated with health and diseases. Sweeteners are widely used food additives, although many studies using animal models have linked sweetener consumption to gut microbial changes and health issues. Whether sweeteners directly change the human gut microbiome functionality remains largely unknown. In this study, we systematically investigated the responses of five human gut microbiomes to 21 common sweeteners, using an approach combining high-throughput *ex vivo* microbiome culturing and metaproteomics to quantify functional changes in different taxa. Hierarchical clustering based on metaproteomic responses of individual microbiomes resulted in two clusters. The first cluster was composed of non-caloric artificial sweeteners (NAS) and two sugar alcohols with shorter carbon backbones (4-5 carbon atoms), and the second cluster was composed of sugar alcohols with longer carbon backbones. The metaproteomic functional responses of the second cluster were similar to the prebiotic fructooligosaccharides and kestose, indicating that these sugar alcohol-type sweeteners have potential prebiotic functions. This study provides a comprehensive evaluation of the direct effects of commonly used sweeteners on the functions of the human gut microbiome using a functional metaproteomics approach, improving our understanding of the roles of sweeteners on microbiome-associated human health and disease issues.

## Introduction

Dietary components, which include carbohydrates, proteins, fats, minerals, food additives, and other compounds, have been shown to play major roles in shaping the composition and function of the gut microbiome, and the associated health consequences.^1,2^ Sweeteners are food additives used to increase the sweetness of food while contributing a negligible amount of energy. In the United States, 25% of children and 41% of adults consumed sweeteners based on data collected from 2009 to 2012,^3^ and the popularity of sweeteners has been continuously increasing.^4^ Sweeteners have been recommended as sugar replacements for better caloric and glycemic control while preserving sweetness.^5^ Sweeteners can be categorized into non-caloric artificial sweeteners (NAS) of high-sweetness intensity, which carry little calories, and sugar alcohols (nutritive sweeteners) with comparable sweetness to sucrose, but indigestible by humans and thus contributing few calories.^6^

Despite the proposed health benefits of sweeteners, many studies have linked their consumption with the development of diseases and metabolic syndrome,^7–9^ some of which were initially intended to be prevented by the use of sweeteners. Sweeteners have been found to induce changes in the gut microbiome composition in animals and humans by metagenomics.^10–12^ Suez *et al*. demonstrated saccharin-induced glucose intolerance in healthy volunteers, and that these effects were transferable to germ-free mice through fecal transplantation.^11^ To date, there have been no systematic studies on sweeteners for their effects on the microbiome Most of the studies were conducted using animal models and only focused on a few sweeteners,^11,13,14^ and comparison across small-scale studies is challenging due to the variation of experimental approaches.

Here we report a systematic study of the effects of sweeteners on individual human gut microbiomes. Briefly, 21 common sweeteners covering all sweeteners approved by Health Canada (HC),^15^ the U.S. Food and Drug Administration (FDA),^16^ and the European Food Safety Authority (EFSA)^17^ as food additives were tested for their impact on the composition and function of five healthy adult microbiomes using the Rapid Assay of Individual Microbiome (RapidAIM)^18^ consisting of *ex vivo* culturing and metaproteomics analysis. To the best of our knowledge, this is the first study using metaproteomic approaches to simultaneously examine the effects of 21 sweeteners on human gut microbiomes. Our results revealed that the sweetener-induced metaproteomic responses of individual microbiomes had two major patterns which were associated with the chemical properties of the sweeteners.

## Results

### Functional profiles of individual microbiomes were altered by sweeteners

In this study, a total of 197 samples (including quality controls and technical replicates) were analyzed by LC-MS/MS. The average MS/MS identification rate was 33.8 ± 6.7%. An average of 8,332 ± 1,744 peptides were identified, and 3,428 ± 533 protein groups were quantified from each sample. Comparison of Bray-Curtis distances between the sweetener-treated microbiome and their non-treated microbiome’s metaproteomics profiles revealed seven sweeteners (xylitol (XYL), isomalt (ISO), maltitol (MAL), lactitol (LAC), sorbitol (SOR), hydrogenated starch hydrolysis (HSH), and mannitol (MAN)) that significantly altered the metaproteome across all five individual microbiomes (Figure 1B). Out of the eight tested sugar alcohols, only erythritol (ERY) did not show significantly altered metaproteomes. PCA of all samples based on protein groups LFQ intensities showed, as expected, strong inter-individual variations (Figure 1C). This was due to the nature of the mixture distribution of each protein group among different individuals. We used an empirical Bayesian algorithm^19^ to fit each mixture distribution to an empirical distribution so as to reduce the effect of individual variance on the dataset. PCA of the data following transformation showed that the control samples of different individuals now clustered together (Figure 1D). In agreement, the sweeteners mentioned above, and controls showed better separation than other groups (Figure 1E and Supplementary Figure S3). In addition, monk-fruit extract (MFE)-treated microbiomes also showed a distinct cluster away from the PBS group (Supplementary Figure S3).

**Figure 1.**
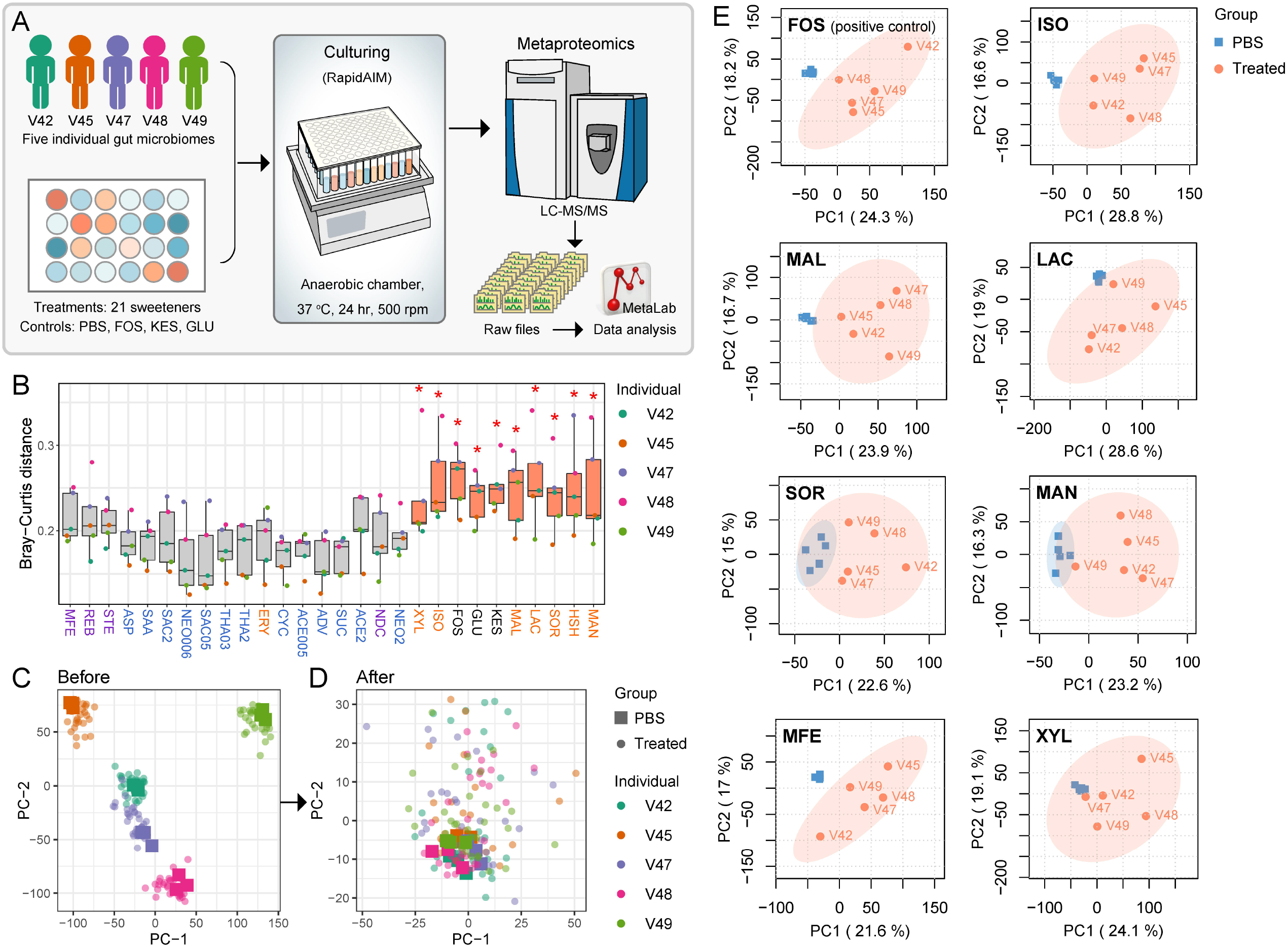
Sweeteners induce metaproteomic changes in the individual microbiomes. Twenty-one sweeteners were analyzed in the study, including four sweeteners tested at two different concentrations (2mg/mL and cADI). Each sweetener is represented by a three-letter abbreviation. SAC2, NEO2, THA2, and ACE2 correspond to sweeteners at 2 mg/mL. SAC05, NEO006, THA03, and ACE005, correspond to sweeteners at cADI (see Supplementary Table S1 for abbreviations and specific concentrations). (A) Workflow combining *ex vivo* culturing and metaproteomics to study the effect of common sweeteners on the gut microbiome. (B) Bray-Curtis distance of protein groups LFQ intensities between sweetener-treated groups and PBS control for each microbiome. Boxes span the interquartile range; jitter colors indicate microbiome number. *p < 0.05, Wilcoxon rank sum test between each group and the average distance among control sample triplicates. Colors of sweetener abbreviations: orange – sugar alcohols, purple – glycoside-type NAS, blue – other NAS, black – controls. (C) PCA score plot generated from protein groups LFQ intensities of all samples (D) PCA score plot after Combat transformation to remove inter-individual variances. (E) Individual PCA score plots of microbiomes treated with positive control FOS and a subset of sweeteners showing separation from the PBS control (based on data after empirical Bayesian transformation).

### Sweeteners induce taxonomic changes in microbiomes

We evaluated the effect of sweeteners on total microbial biomass by measuring the total proteins obtained in each sample using DC assay as described in Section 2.3 (Supplementary Figure S4). For most sweeteners, their effect on the individual microbiomes varied, while ISO and thaumatin at 2 mg/ml (THA2) increased the total biomass in all five microbiomes.

Genus-level protein abundance was then calculated from the distinctive peptide intensities of each genus measured by LC-MS/MS and the total microbial protein biomass of each sample. Most sweeteners showed significant effects on the abundance of at least one genus (Figure 2A). Clustering based on effects of genus-level biomass shows that the microbial community composition was affected in patterns by different classes of sweeteners (Figure 2B). Glycoside-type NAS rebaudioside A (REB), neohesperidin dihydrochalcone (NDC), and MFE, as well as sugar alcohols MAL, LAC, XYL, and ISO clustered together and formed a distinct cluster from other sweeteners and controls. This cluster showed significant increases in several genera such as *Dialister, Parabacteroides, Ruminococcus, Phascolarctobacterium, Butyrivibrio, Blautia*, and *Marvinbryantia*. Sugar alcohols SOR, MAN, and HSH clustered with controls FOS, KES, and GLU. This cluster was featured with significant increases in Actinobacteria genera *Bifidobacterium* and *Collinsella*, and decreases in *Dorea, Clostridium, Lachnoclostridium, Alistipes, Roseburia*, and *Flavonifractor*. Genera *Coprococcus, Oscillibater, Anaerostipes*, and *Butyrivibrio* were increased by XYL, but not by any other sugar alcohols.

**Figure 2.**
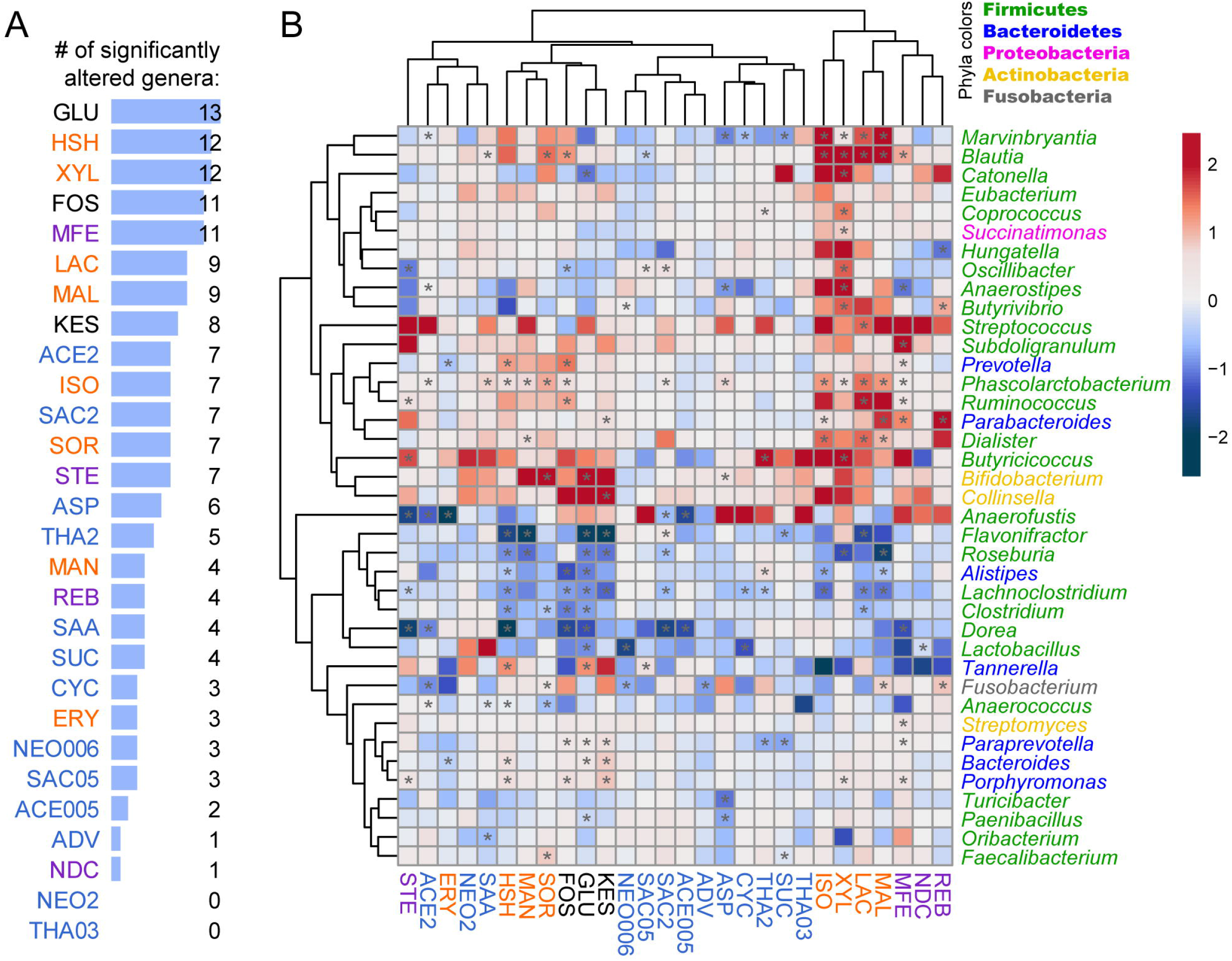
Sweeteners induced gut microbiome taxonomic changes. Sweeteners are named as in Figure 1. (A)The number of the significantly affected genus by each sweetener. (B)Heatmap showing log_2_ fold change of genus-level protein abundance of sweetener-treated samples versus PBS control. For each treatment, the averaged genus biomass of all five microbiomes were used for coloring and clustering. *p < 0.05, Wilcoxon rank sum test. Genera that were detected in PBS controls in at least four out of the five microbiomes are shown.

### Functional metaproteomics segregates the sweeteners into two groups

Of the identified protein groups, 93.5% had COG functional annotation. Sweeteners were categorized into two major clusters using COG abundances (Figure 3A and Supplementary Figure S5). Bootstrapping of the two major clusters gave scores of 0.983 and 0.956, respectively, indicating high clustering robustness (Figure 3A). We named the two clusters as the “NAS” and the “CHO” cluster according to the properties of sweeteners. Statistical analyses at the COG category level identified 14 out of 21 COG categories significantly altered by at least two compounds (Figure 3 B-G and Supplementary Figure S5).

**Figure 3.**
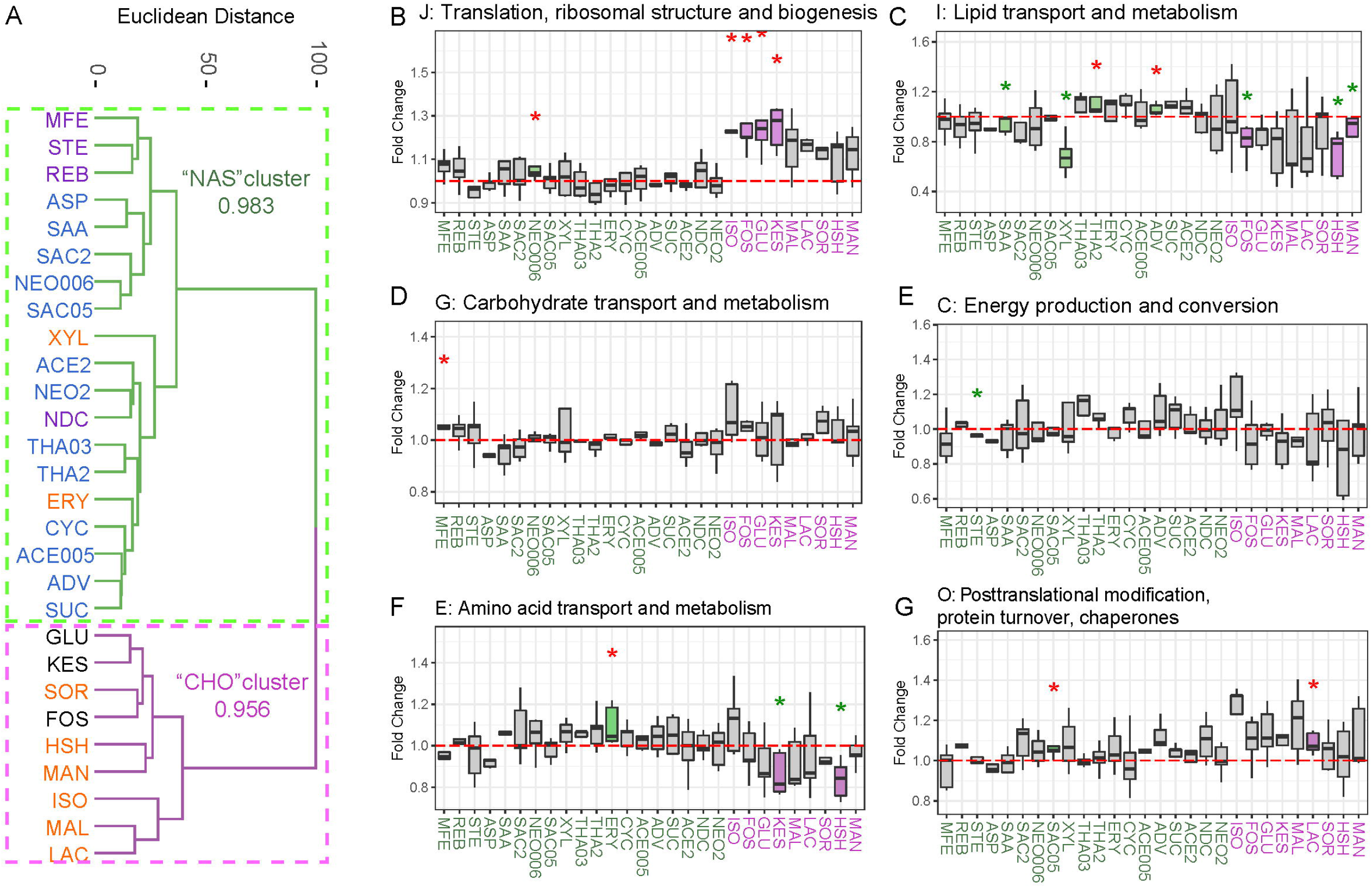
Sweeteners induced functional changes in the gut microbiome. Saweeteners are named as in Figure 1. (A) Clustering of sweeteners based on induced functional responses. Euclidean distances are calculated between sweeteners based on averaged log_2_ fold change of COG abundances of sweetener-treated samples versus PBS control. Bootstrapping scores of the two major clusters are shown. (B) – (G) Fold change between the treated group and PBS control of several COG categories. Colored boxes indicate significantly changed COG categories. Red and green asterisks indicate significant increase and decrease, respectively. *p < 0.05, Wilcoxon rank sum test. Responses of all other COG categories are shown in Supplementary Figure S5.

In the “NAS” cluster, all NASs were included, plus sugar alcohols ERY and XYL. ERY and XYL, have shorter carbon backbones, with ERY backbone comprising 4 carbon atoms and XYL comprising 5 carbon atoms (Supplementary Figure S1). Interestingly, although four sugar alcohols MAL, LAC, XYL, and ISO were clustered with three of the NAS - REB, NDC, and MFE - in the taxonomic analysis in Figure 2, the functional responses of the microbiome to these two clusters were different. While XYL still belonged to the “NAS” cluster, three other sugar alcohols (i.e., MAL, LAC, and ISO) belonged to the “CHO” cluster. XYL showed marked effects on the metaproteome, including significantly decreased lipid transport and metabolism, cell motility, and significantly increased coenzyme transport and metabolism (Figure 3C and Supplementary Figure S5). Extracellular structures were significantly promoted by the “MFE, STE, REB” sub-cluster (Supplementary Figure S5). The “CHO” cluster included all remaining sugar alcohols and positive controls, all of which are carbohydrates. In the “CHO” cluster, sugar alcohols SOR, MAN, LAC, MAL, ISO, and HSH cannot be digested by the human body. Therefore, despite incomplete absorption of sugar alcohols in the small intestine,^20^ they can reach the colon intact, serving as substrates for microbial fermentation to produce hydrogen gas, carbon dioxide, methane, and short-chain fatty acids (SCFA).^21^ Compounds in this cluster were found to induce marked responses in the metaproteomes in a similar pattern, including an increase of translation, ribosomal structure, and biogenesis and a decrease of lipid transport and metabolism (Figure 3B and C). In addition, proteins with only general function prediction in the COG database were also significantly increased by a subset of the CHO cluster (Supplementary Figure S5). Sugar alcohols are shown to induce gastrointestinal symptoms including bloating, laxative effect, and abdominal pain.^22^ Accordingly, we showed that some sugar alcohols, such as ISO, significantly reduced cell motility, which has been reported to be associated with increased susceptibility to intestinal expulsion and larger fluctuation in absolute abundance.^23^

### “CHO” cluster sweeteners had prebiotic-like effects on Clostridia

Partial Least Square Discriminant Analysis (PLS-DA) was performed to identify the most important differences in functional effects between the “CHO” cluster and “NAS” cluster (Figure 4A and 4B). 214 of 3,608 protein groups had a VIP score > 2 in the first five components, which indicates that these proteins explain the most important differences in functional effects between the two clusters. These proteins were referred to as discriminative proteins. The intensity profiles of these 214 discriminative proteins in response to the treatment of different sweeteners segregated into two well-defined groups (Figure 4C). 106 of the discriminative proteins that had a higher abundance in the “CHO” cluster were referred to as CHO elevated group. As sweeteners of the “NAS” cluster were clustered with the PBS group, indicating that proteins enriched in the “NAS” cluster were decreased in the “CHO” cluster, 108 of the discriminative proteins had a higher abundance in the “NAS” cluster were referred as CHO depleted group.

**Figure 4.**
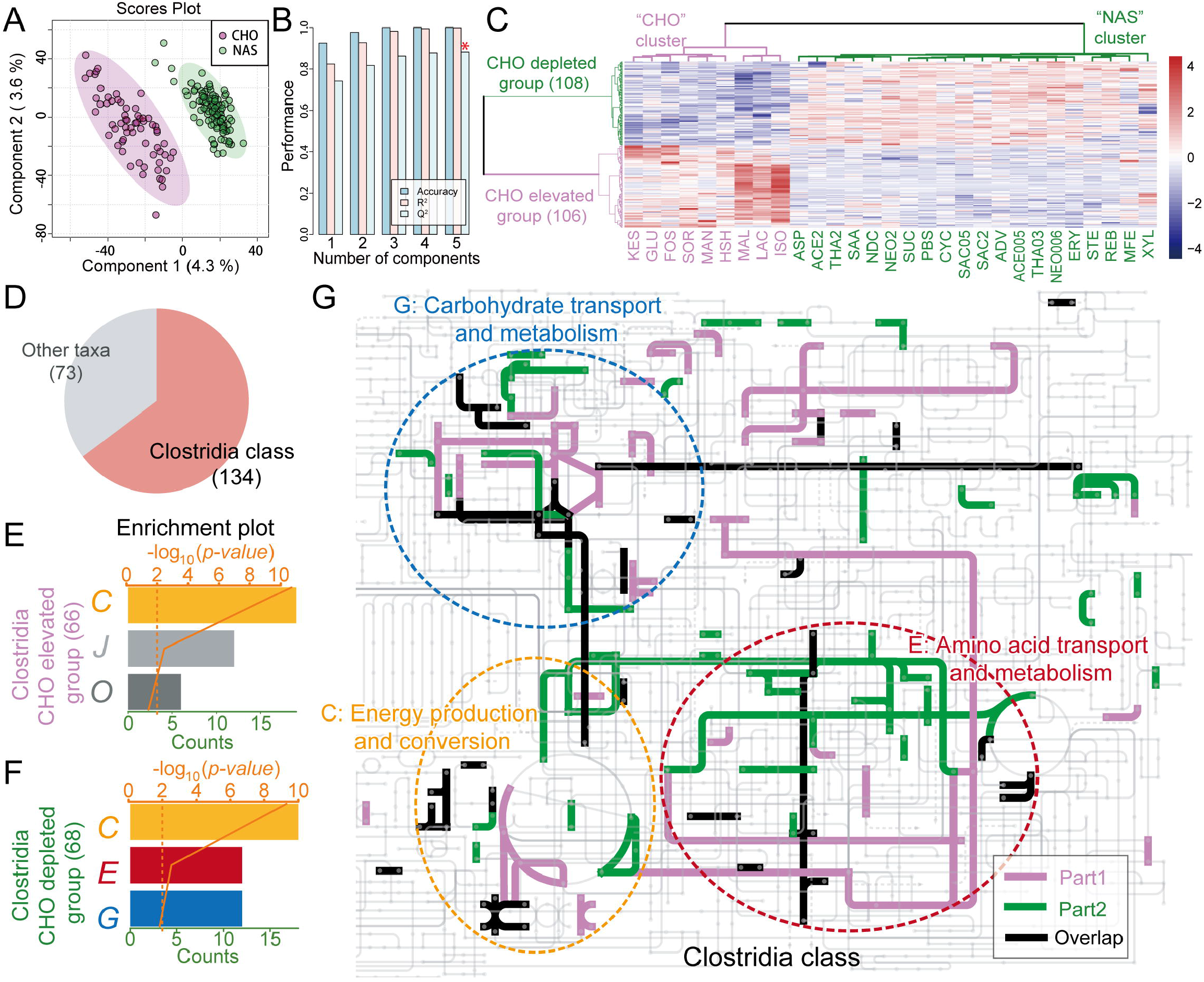
Taxonomy and functional profiles of discriminative proteins for the “CHO” cluster and “NAS” cluster revealed by PLS-DA. (A) PLS-DA scores plot for differential protein profiles across the “CHO” cluster and “NAS” cluster sweeteners. (B) PLS-DA cross-validation results. (C) Heatmap of the intensity of 214 discriminative proteins under the treatment of different sweeteners (sweeteners are named as in Figure 1). (D) The taxonomy sources of discriminative proteins. (E) and (F) Enriched COG categories of discriminative proteins in Clostridia. (G) Pathways of CHO elevated group and CHO depleted group discriminative proteins in Clostridia. Proteins related to COG Categories C, G, and E were framed in dash circles.

As not all the identified peptides had taxonomy information from the Metalab, two hundred and seven out of 214 discriminative proteins were assigned taxonomy information from peptides as described in the methods, and 62.6% of the discriminative proteins were from class Clostridia (Figure 4D). Functional enrichment analysis of each group of discriminative proteins from Clostridia was performed. CHO elevated group was enriched in COG categories C, J, and O, while CHO depleted group was enriched in COG categories C, E, and G (Figure 4E and 4F). Although both the CHO increased group and CHO depleted group from Clostridia were enriched in COG Category C, the enriched proteins from the two groups had different functions (Supplementary Table S2). Pathway analysis also showed that proteins from two groups were from different metabolic pathways (Figure 4G). In particular, iron-dependent proteins that are involved in the TCA cycle between succinyl-CoA and Malate, and its linkage to oxidative phosphorylation (COG0427, COG0479, COG1145, COG1838, COG1951, COG2048) were highly enriched. The “CHO” cluster sweeteners had similar effects on the protein abundance of discriminative proteins with FOS and KES which are commonly used prebiotics.

## Discussion

Although there have been studies on the effects of sweeteners on the human gut microbiome,^11,12^ the comparison of the effects of a large number of different sweeteners on human gut microbes has not been reported. In this study, we investigated the taxonomic and functional responses of five individual human microbiomes to 21 common sweeteners. Among the 21 sweeteners, thirteen are NAS with diverse chemical properties and high sweetness intensities, and eight are sugar alcohols, which are carbohydrates with low digestibility.^22^

We observed that seven sweeteners (XYL, ISO, MAL, LAC, SOR, HSH, MAN) significantly altered the metaproteome across the five gut microbiomes (Figure 1B). The remaining sweeteners had no global metaproteome effects. Although MFE showed a clear separation between control and treatment by PCA, the alteration was not significant using Bray-Curtis distance. A previous study reported that saccharin (SAC) alters the gut microbiota and induces glucose intolerance in mice model.^11^ While we did not observe any global effects of saccharin on the metaproteome using our *ex vivo* assay, we observed changes at the individual level indicating an individual-specific response to saccharin. A similar individual-specific response was also observed for sucralose (SUC), which has been shown by others to alter the composition of the gut microbiota in a rat model.^13,14^ Although several sweeteners significantly changed the metaproteome of the gut microbiome, most sweeteners had limited effects on the biomass of the microbiome. Only ISO and thaumatin at 2 mg/ml (THA2) led to a biomass increase in all five microbiomes (Supplementary Figure S2).

The genus-level taxonomic changes of the gut microbiome under different sweeteners were evaluated. A sweetener cluster consisting of both glycoside-type NAS (REB, NDC, MFE) and sugar alcohols (MAL, LAC, XYL, ISO) significantly increased protein abundance from several genera in Firmicutes. A single species-based study revealed XYL was largely utilized by *Anaerostipes caccae* from Firmicutes to produce butyrate and promoted the growth of the species.^24^ The effects of sweeteners on other genera from Firmicutes need to be further studied. In addition, controls FOS, KES and GLU were clustered with sugar alcohols SOR, MAN, and HSH. It is noted that this clustering result was different from the functional profile clustering. While metaproteomics measures the abundances of different taxa by summarizing the overall protein intensities, functional profiles provided a deeper layer of information by comparing functional proteins in the community and are more relevant to the actual functionality and state of the microbiomes.

By analyzing the functional profile of the cultured microbiomes, we segregated all the sweeteners into two clusters, “NAS” and “CHO”. To further investigate the different effects of two clusters of sweeteners on the human gut microbiome, discriminative proteins with PLS-DA VIP > 2 were identified. FOS and KES clustered with the “CHO” cluster, which indicated that the “CHO” cluster might have similar functional effects with FOS and KES. Most of the discriminative proteins were from Clostridia. Clostridia accounts for at least 10-40% of the total bacteria in gut microbiota^25^ and members from this class have significant potential as probiotics^26^.

Functional enrichment analysis of discriminative proteins from Clostridia revealed that proteins with higher abundances in the “CHO” cluster (CHO elevated group) and proteins with higher abundances in the “NAS” cluster (CHO depleted group) were both enriched in COG category C. Enriched proteins from both groups corresponding to different functions (Supplementary Table S2), with differences in specific pathways as shown by the metabolic pathway analysis (Figure 4G). In particular, the enrichment of iron-dependent pathways in TCA cycle and oxidative phosphorylation indicated increased cellular energy metabolism in Clostridia in response to the “CHO” cluster. Other proteins from the two groups also corresponded to different pathways indicating that the two clusters of sweeteners had effects on the different metabolic pathways of Clostridia. In addition, CHO elevated group was enriched in COG function categories J (Translation, ribosomal structure and biogenesis) and O (Post-translational modification, protein turnover, chaperone functions). The higher abundance of proteins from these two categories indicated that the “CHO” clustered sweeteners might promote the cell division of Clostridia. A previous study showed that traditional prebiotics “FOS” and probiotics *Lactobacillus* can increase the proportion of Clostridia in gut microbiomes,^27^ and “CHO” clustered sweeteners might have similar potential with these prebiotics and probiotics. Discriminative proteins with higher abundance in the “NAS” cluster (CHO depleted group) were enriched in COG categories C, G, and E (Amino acid transport and metabolism). The functional annotation of enriched proteins (Supplementary Table S2) showed that a certain amount of these discriminative proteins in COG categories C (5 of 18) and G (4 of 12) were related to glycerol transport and alcohol metabolism. This suggested inhibition of the glycerol transport and alcohol metabolic pathways by the “CHO” cluster.

## Conclusion

We examined the effect of 21 sweeteners on the *ex vivo* human gut microbiome from five individuals. Seven sweeteners significantly altered the metaproteomes across the five microbiomes. Functional profile of the microbiomes clustered all the sweeteners into the “NAS” cluster and the “CHO” cluster. Prebiotics FOS and KES clustered with the “CHO” cluster. Most discriminative proteins for the two clusters were from Clostridia. Functional enrichment analysis and pathway analysis revealed that two cluster sweeteners had effects on the different pathways of Clostridia. “CHO” cluster sweeteners had similar effects as the prebiotics FOS and KES. Our study revealed the functional effects of sweeteners on the human microbiome and suggested the prebiotic potential of the sugar alcohol sweeteners.

## Materials and Methods

### Sweeteners and the determination of concentrations

The concentration of sweeteners used in the assay (Supplementary Figure S1) was determined based on their consumption levels in the general public, the acceptable daily intake (ADI), and the proportion of consumed sweeteners that could reach the colon (Supplementary Figure S2). Twelve of the tested sweeteners had ADI data defined by FDA or EFSA. The culturing concentration that met the ADI (cADI) for each sweetener was calculated based on consumption from an average participant body weight of 70.3 kg normalized to 200 g of colon content. And the calculation of the cADI was conducted based on the following formula:

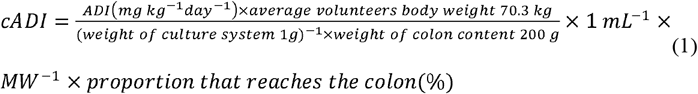

Sweeteners with cADI values much lower than 2 mg/mL were tested both at cADI and at 2 mg/mL to facilitate the comparison of their effects on the microbiome with other sweeteners. For sweeteners without defined ADI, in the case of all sugar alcohols, the concentration used was standardized to 2 mg/mL, which would represent consumption levels lower than those in the general public.^28–30^ Since the cADI of advantame exceeded its solubility and the ADI would represent a consumption level much higher than that in the general public,^31^ advantame was tested at 2 mg/mL, about 1/5 of cADI. In addition aspartame (ASP), thaumatin (THA), and salt of aspartame-acesulfame (SAA) are known to be completely metabolized before reaching the colon,^32,33^ resulting in cADI of 0. A study has shown that carbohydrates that can be absorbed by the small intestine can still enter the large intestine and be fermented by colonic microbiota.^34^ We included ASP, THA, and SAA in this study, assuming a portion of these sweeteners may also reach the colon.

### Human stool sample collection and culturing

The study was approved by the Ottawa Health Science Network Research Ethics Board at the Ottawa Hospital (# 20160585-01 H). Stool samples from five healthy volunteers (from 23 to 48 years old, 3 males and 2 females) were included in this study. Exclusion criteria included the diagnosis of irritable bowel syndrome (IBS), inflammatory bowel disease (IBD) or diabetes; antibiotic use or gastroenteritis episode in the last three months; use of pro-/pre-biotic, laxative, or antidiarrheal drugs in the last month; or pregnancy. Volunteers included in this study were self-accessed as non-sweetener consumers. The stool was collected on-site and immediately transferred into an anaerobic workstation (5% H_2_, 5% CO_2_, and 90% N_2_ at 37°C). A 20% (w/v) stool slurry was made in sterile pre-reduced PBS (pH 7.6) containing 10% (v/v) glycerol and 1 g/L L-cysteine, vortex homogenized and filtered through gauze. The filtered slurry aliquots were stored at -80°C until culturing.

The frozen fecal samples were thawed at 37°C and immediately transferred into the anaerobic workstation, vortex homogenized, and inoculated at 2% (w/v) into 96 deep-well plates containing pre-reduced, optimized microbiome culture medium^18,35^ and a sweetener (manufacturers and concentrations used shown in Supplementary Table S1), a positive control (2 mg/ml fructooligosaccharide or 2 mg/ml kestose), or the negative control (PBS) in each well. The plates were shaken at 500 rpm on shakers (MS3, IKA, Germany) in the anaerobic workstation at 37°C for 24 hours. Following anaerobic culture, samples were processed to isolate bacterial pellets as previously described.^18^ Bacterial pellets stored at -80°C until LC-MS sample preparation.

### Metaproteomic sample preparation

Proteins were extracted from bacterial pellets as per Li *et al*.^18^ Briefly, bacterial pellets were resuspended in bacterial lysis buffer containing 4% (w/v) sodium dodecyl sulfate (SDS), 8 M urea, 50 mM Tris-HCl (pH 8.0), cOmplete™ mini protease inhibitors, and PhosSTOP inhibitor (MilliporeSigma). Samples were sonicated (Qsonica, USA) at 8°C, 50% amplitude, for 10 min with a 10 s on 10 s off working cycle. Lysates were centrifuged at 16000 g at 8°C for 10 min to remove cell debris. Supernatant protein concentrations were measured with Bio-Rad’s detergent compatible (DC) protein assay reagent (USA) in triplicate and used to calculate the total biomass of the microbiome after culturing.

Proteins from each sample were precipitated overnight at -20°C by mixing lysis supernatant with a protein precipitation buffer containing 50% (v/v) acetone, 50% (v/v) ethanol, and 0.1% (v/v) acetic acid at 1:5 ratio (v/v). Precipitated proteins were collected by centrifugation at 16000g at 4°C for 25 min. Proteins were washed with 1ml -20°C acetone three times and pelleted by centrifugation at 16000g at 4°C for 25 min. Following acetone washes, proteins were dissolved in 6 M urea and 50 mM ammonium bicarbonate (ABC, pH 8.0). Protein concentrations were determined as above. 50 ug protein aliquots were reduced with 10mM dithiothreitol at 56 °C for 30 min with shaking. Protein alkylation was then performed at 20 mM iodoacetamide in the dark for 40 min at room temperature. Samples were diluted 10-fold with 50 mM ABC (pH 8.0). Trypsin (Worthington Biochemical Corp., Lakewood, NJ) was added to a mass ratio of trypsin: protein = 1:50, and digestion was performed at 850 rpm at 37 °C on for 19 hours. Trypsin digestion was stopped by adding 50 μL 10% (v/v) formic acid to the final pH of 2 to 3. Desalting was performed on in-house made 96-channel filter tip plates packed with 5 mg 10-μm C18 resin (Dr. Maisch GmbH, Ammerbuch, Germany). Desalted samples were freeze-dried and stored at -20°C.

### HPLC-MSMS analysis

An Eksigent nano-LC system (nano2D ultra) coupled with a Q Exactive mass spectrometer (Thermo Fisher Scientific Inc.) was used for analysis. Tryptic peptides were reconstituted in 0.1% (v/v) formic acid to approximately 0.25 μg/μL. 1 μg of peptides were loaded. The column used for peptide separation was of 75 μm inner diameter and 15 cm long, packed with reverse phase C18 resin (1.9μm/120 Å ReproSil-Pur C18 resin, Dr. Maisch GmbH, Ammerbuch, Germany). A 90-min gradient was used with acetonitrile changing from 5% to 30% (v/v) at a flow rate of 300 nL/min. Solvent A was composed of 0.1% (v/v) formic acid, solvent B was composed of 0.1% (v/v) formic acid and 80% v/v acetonitrile. MS analysis was performed with a Q Exactive mass spectrometer (ThermoFisher Scientific Inc.). Full MS scans were performed from 300 to 1800 m/z, data-dependent MS/MS scans were performed for the 12 most intense ions. MS and MS/MS scans were performed with resolutions of 70000 and 17500, respectively. Samples were loaded in a randomized order. In this study, 197 samples were analyzed over a period of 23 days. All raw data from LC-MS/MS have been deposited with the ProteomeXchange Consortium (http://www.proteomexchange.org) via the PRIDE^36^ partner repository via the dataset identifier PXD030458.

### Protein identification, quantification, and profile

The Metalab software (version 1.2.0) was used for peptide/protein identification and quantification, peptide taxonomy assignment, and protein functional annotation.^37^ The searches were performed against a database based on the integrated gene catalog (IGC) which included close-to-complete sets of genes for most gut microbes.^38^ Carbamidomethyl (C) was set as a fixed modification, and protein N-terminal acetylation (protein N-term) and Oxidation (M) were set as variable modifications.

Analysis of changes in the gut metaproteome was based on the quantified protein groups. Label-free quantification (LFQ) intensities of each protein group were first normalized by the estimated size factor calculated using the R package “DESq2”.^39^ Bray-Curtis distances between samples were calculated based on the normalized intensities using the R package “vegan”.^40^ For principal component analysis (PCA), protein groups were filtered based on criteria that the protein group appears in at least one treatment in at least three out of the five tested microbiomes (60%). The intensities were then log_10_-transformed and PCA was performed using the R package “stats”. To reduce the effect of inter-individual microbiome variation on data analysis, the distribution of each protein group among individual microbiomes were fitted to the same empirical distribution using an empirical Bayesian algorithm with ComBat^19^ on iMetalab.ca.^41^

### Microbial taxonomic analysis

Taxonomy assignment of each peptide was performed based on the lowest common ancestor (LCA), with the abundance of each taxon calculated by summarizing the intensities of all distinctive peptides assigned to this taxon. Relative abundance of taxa on a specific taxonomic rank was calculated by normalizing to the summed abundance of all taxa on this rank. For comparison of absolute taxa abundance between samples, relative abundances were multiplied by the total microbial biomass calculated using protein concentration data. Fold changes were calculated between sweetener-treated samples and PBS control from the same microbiome.

### Microbial function analysis

Functional annotation was carried out in the Metalab software, and each identified protein group was assigned to a cluster of orthologous groups (COG). The relative abundance of each COG was calculated based on the summed LFQ intensities of all assigned protein groups. Clustering of sweeteners was based on the fold change of relative COG intensities, averaged across all tested sweeteners. The Euclidean distance between sweeteners was calculated and the clustering was performed with the “ward.D” method, using the R package “stats”. Bootstrapping evaluation^42^ of the two major clusters was performed using the R package “fpc”^43^ with the number of resampling runs being 100.

Partial least squares discriminant analysis (PLS-DA) was performed to identify discriminative proteins that reveal the difference in the effects of the two clusters of sweeteners. PLS-DA was performed in MetaboAnalyst 5.0^44^ and discriminative proteins were selected with a VIP (Variable Importance in Projection) > 2. The taxonomy sources of the discriminative proteins were obtained by the method described in a previous study.^45^ Briefly, among all the peptides contained in the discriminative protein, the taxonomy source of the peptide with the most detailed taxonomy information was regarded as the taxonomy source of the protein. All the peptides from a discriminative protein were acquired from the protein groups table from the MetaLab output, and the taxonomy information of the peptides was obtained through the taxonomy table from Metalab. Functional enrichment analysis of these discriminative proteins was performed using an online enrichment analysis tool (https://shiny.imetalab.ca/), p-value threshold was set as < 0.05. Visualization of pathways was performed using COG accession numbers in iPath3.^46^

## Data Accessibility

All raw data from LC-MS/MS have been deposited with the ProteomeXchange Consortium (http://www.proteomexchange.org) via the PRIDE partner repository (PXD030458).

## Author Contributions

D.F. and W.W. designed the study. J.M. coordinated sample collection and biobanking. W.W., J.M., L.L., and K.W. performed the experiments. Z.S, W.W., L.L., X.Z., and Z.N. performed data analysis. Z.S, W.W., L.L., X.Z., Z.N., and D.F. wrote the manuscript. All authors participated in the data interpretation, discussion, and edits of the manuscript

## Institutional Review Board Statement

The study was approved by the Ottawa Health Science Network Research Ethics Board at the Ottawa Hospital (# 20160585-01 H).

## Informed Consent Statement

Informed consent was obtained from all subjects involved in the study.

