## Supplementary Materials for "Comprehensive assessment of functional effects of commonly used sweeteners on *ex vivo* human gut microbiome"

### Supplementary Figures

**Supplementary Figure S1.** Structure of sweeteners.

**Supplementary Figure S2.** Procedures used to determine sweetener concentrations.

**Supplementary Figure S3.** Individual PCA plots of other sweeteners, positive control KES and dietary sugar control GLU.

**Supplementary Figure S4.** Fold change of total protein amount obtained for each treatment group compared to PBS control.

**Supplementary Figure S5.** Responses of other COG categories in addition to Figure 3.

### Supplementary Tables

**Supplementary Table S1** Summary of sweeteners.

**Supplementary Table S2** Functional annotation of enriched discriminative proteins from Clostridia.

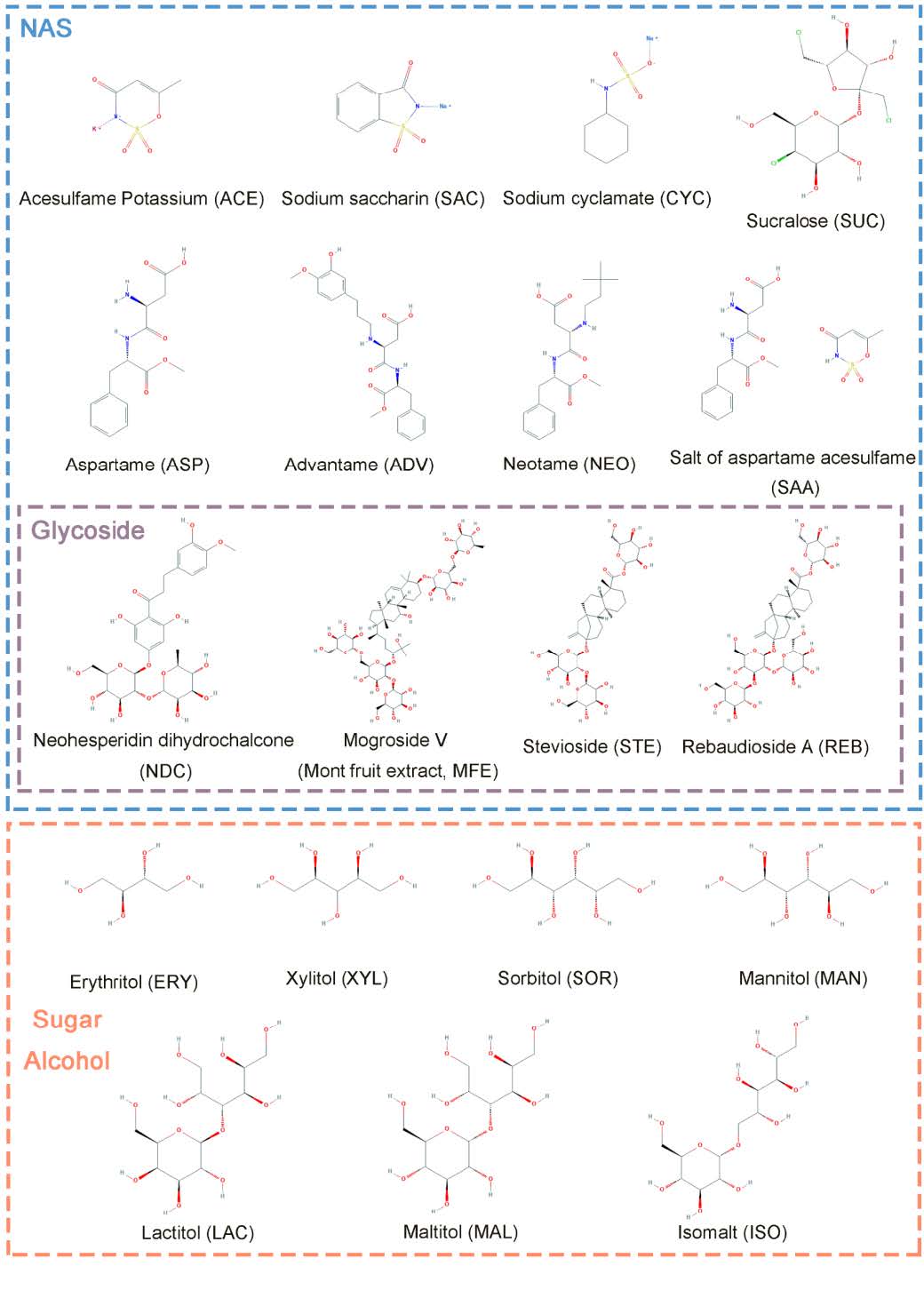
**Supplementary Figure S1.** Structure of sweeteners. Thaumatin (THA) and hydrogenated starch hydrolysates (HSH) were not included in this figure as they are mixtures and have no structural formula.

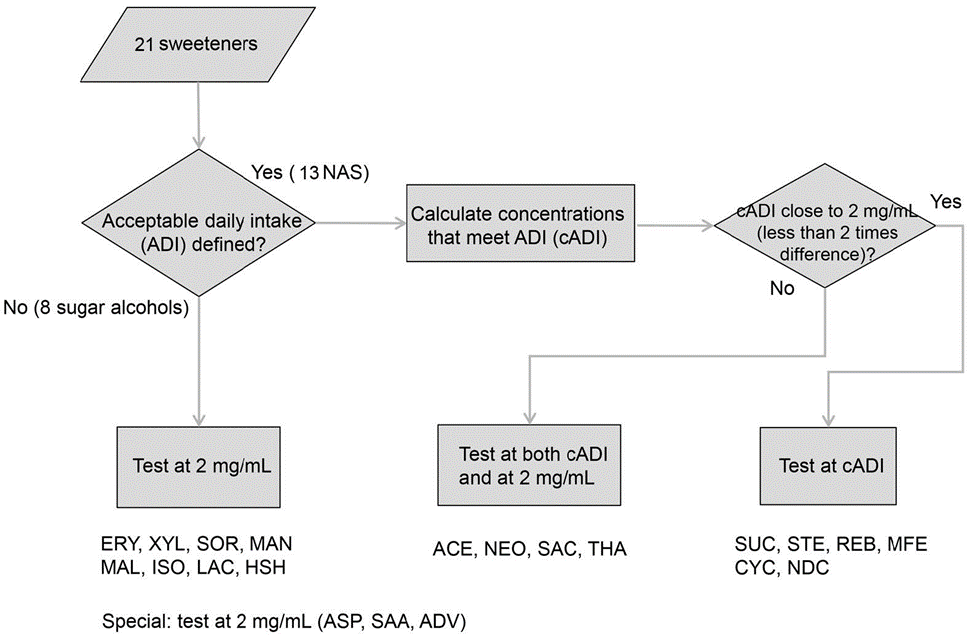
**Supplementary Figure S2.** Procedures used to determine sweetener concentrations.

**Supplementary Figure S3.** Individual PCA plots of other sweeteners, positive control KES and dietary sugar control GLU
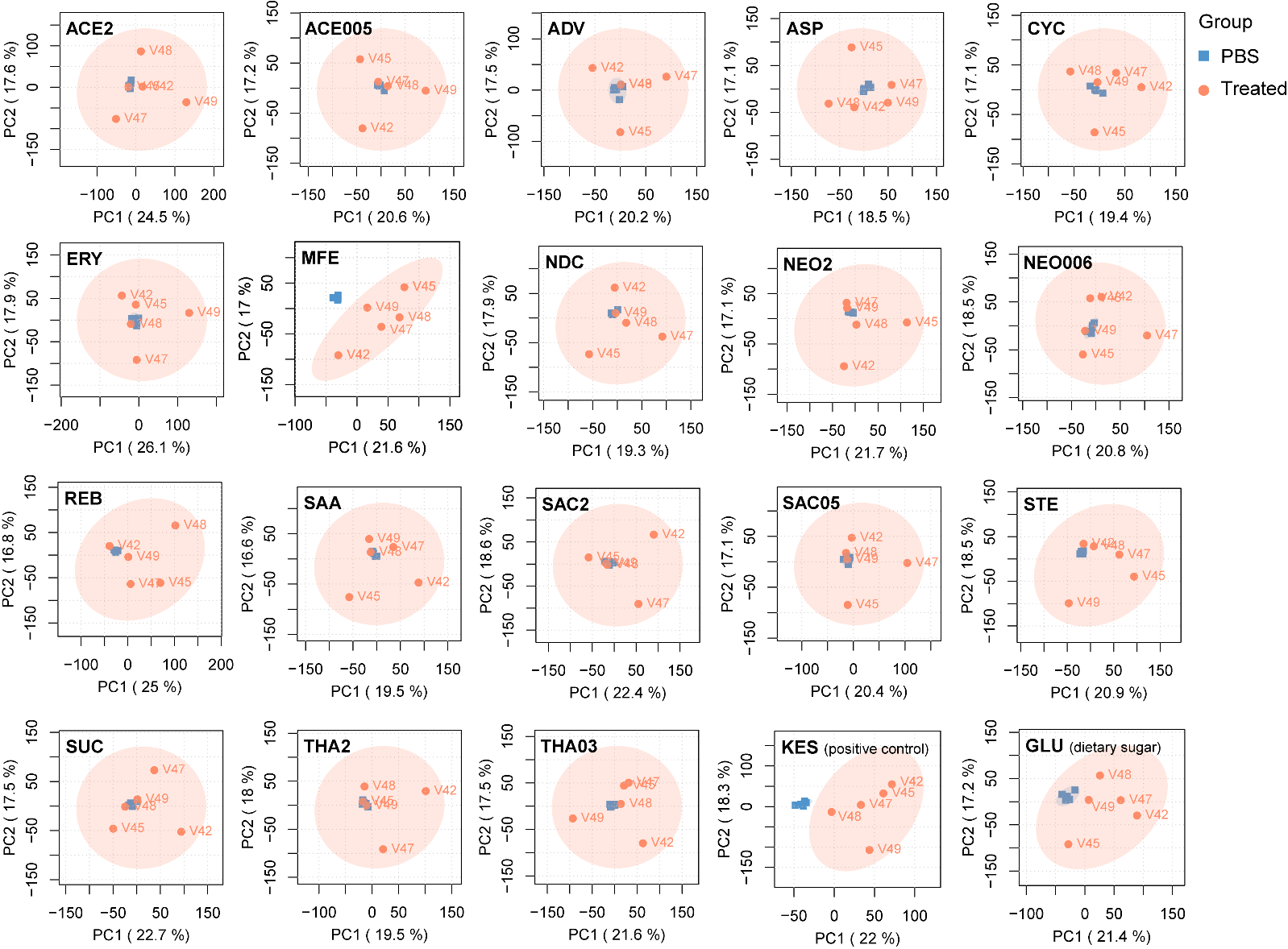
.

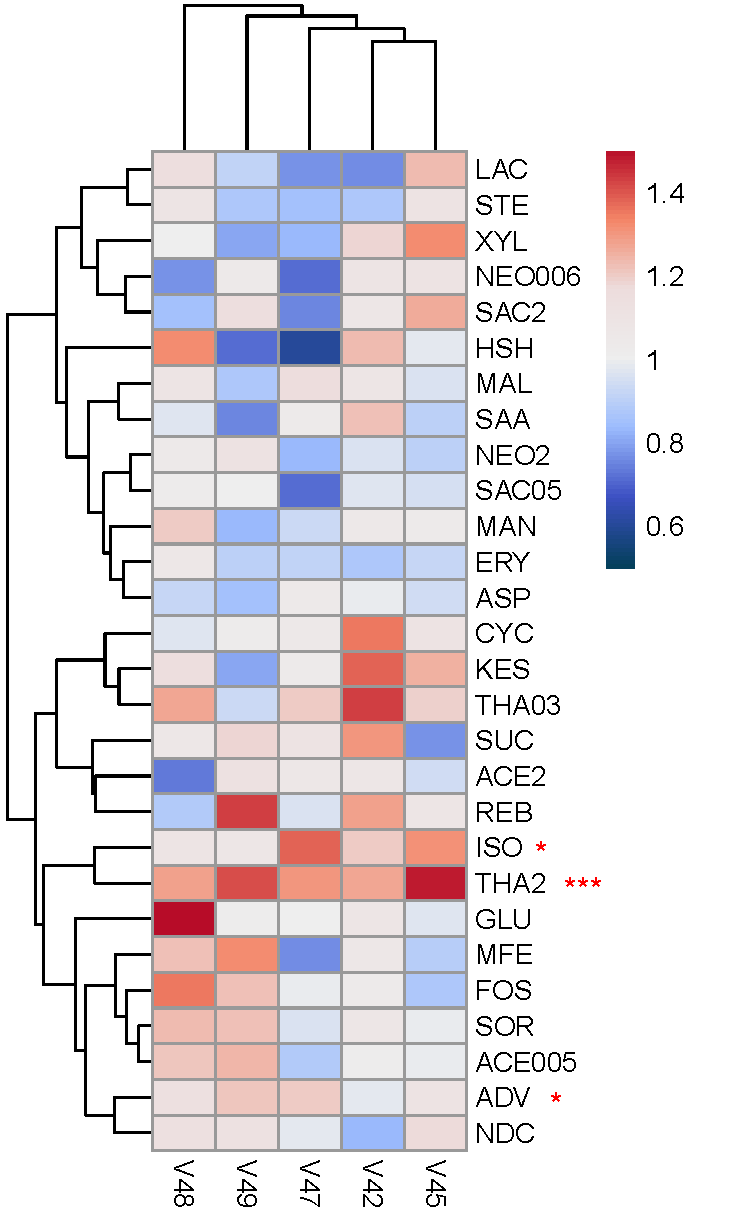
**Supplementary Figure S4.** Fold change of total protein amount obtained for each treatment group compared to PBS control. * and *** denotes p < 0.001 and p < 0.05, respectively, by two-sided t-test.

**Supplementary Figure S5.** Responses of other COG categories in addition to Figure 3
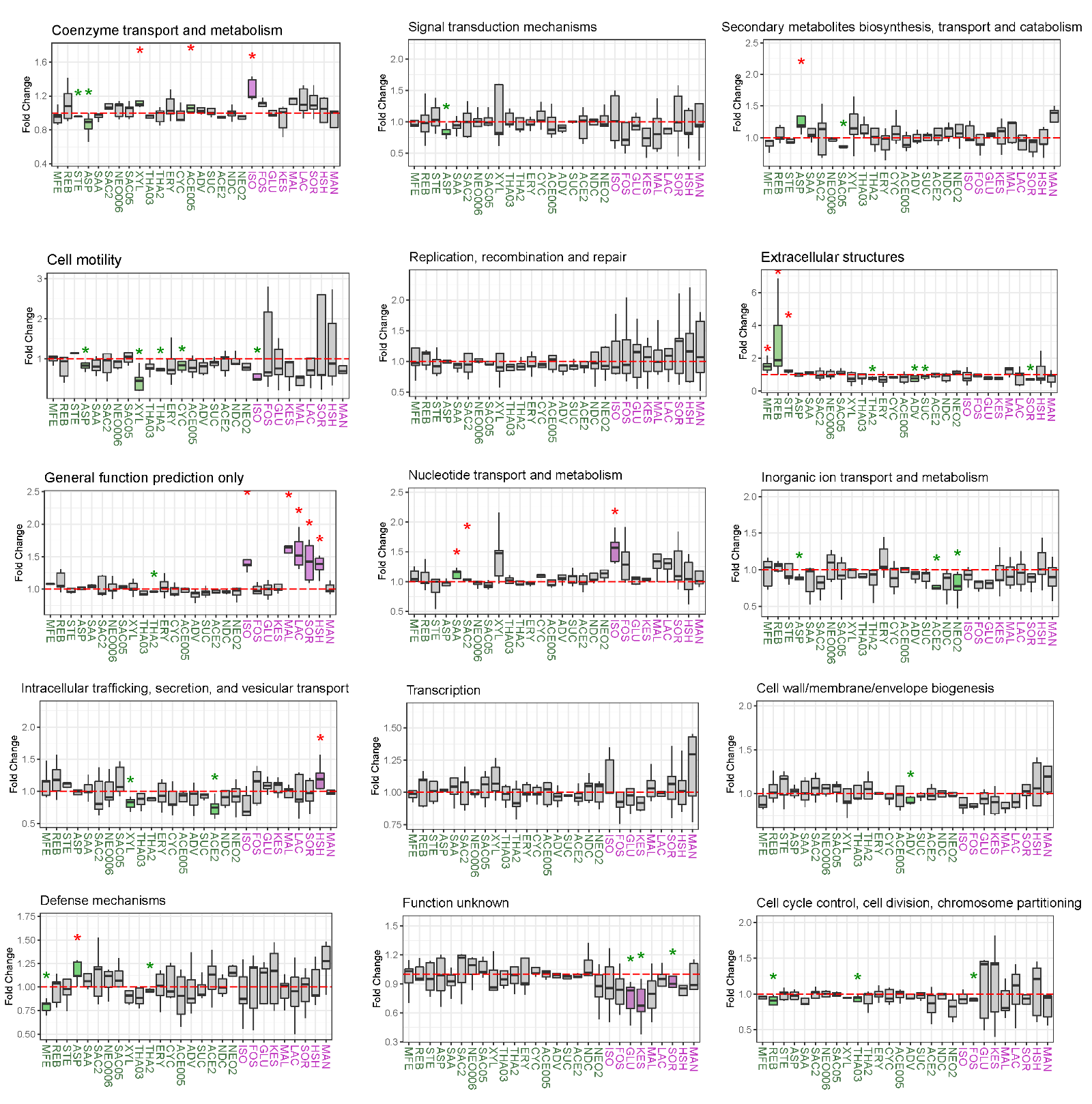
.

**Supplementary Table S1** Summary of sweeteners.

| Sweetener* | Classification | Abbreviation in this paper | PubChem CID | Molecular Weight (g/mol) | Approved by |
| --- | --- | --- | --- | --- | --- |
| Acesulfame K | NAS | ACE | 11074431.00 | 201.24 | HC, EFSA, FDA |
| Aspartame | NAS | ASP | 134601.00 | 294.30 | HC, EFSA, FDA |
| Advantame | NAS | ADV | 10389431.00 | 458.50 | HC, EFSA, FDA |
| Neotame | NAS | NEO | 9810996.00 | 378.50 | HC, EFSA, FDA |
| Saccharin (Sodium saccharin dihydrate) | NAS | SAC | 517320.00 | 241.20 | HC, EFSA, FDA |
| Sucralose | NAS | SUC | 71485.00 | 397.60 | HC, EFSA, FDA |
| Stevia extract (Stevioside) | NAS | STE | 442089.00 | 804.90 | HC, EFSA, FDA |
| Stevia extract (Rebaudioside A) | NAS | REB | 6918840.00 | 967.00 | HC, EFSA, FDA |
| Monk fruit extract | NAS | MFE | N/A | N/A | HC, FDA |
| Thaumatin | NAS | THA | N/A | N/A | HC, EFSA, FDA |
| Cyclamate (Sodium cyclamate) | NAS | CYC | 23665706.00 | 201.22 | EFSA |
| Neohesperidin Dihydrochalcone | NAS | NDC | 30231.00 | 612.60 | EFSA |
| Salt of Aspartame-Acesulfame | NAS | SAA | 25130065.00 | 457.50 | EFSA |
| Sorbitol (D-Sorbitol) | sugar alcohols | SOR | 5780.00 | 182.17 | HC, EFSA, FDA |
| Mannitol (D-Mannitol) | sugar alcohols | MAN | 6251.00 | 182.17 | HC, EFSA, FDA |
| Isomalt | sugar alcohols | ISO | 88735.00 | 344.31 | HC, EFSA, FDA |
| Maltitol | sugar alcohols | MAL | 493591.00 | 344.31 | HC, EFSA, FDA |
| Lactitol (Lactitol monohydrate) | sugar alcohols | LAC | 3067270.00 | 362.33 | HC, EFSA, FDA |
| Xylitol | sugar alcohols | XYL | 6912.00 | 152.15 | HC, EFSA, FDA |
| Erythritol (meso-Erythritol) | sugar alcohols | ERY | 222285.00 | 122.12 | HC, EFSA, FDA |
| Hydrogenated starch hydrolysates | sugar alcohols | HSH | N/A | N/A | HC, EFSA, FDA |
| Glucose (D-Glucose) | dietary sugar (positive control) | GLU | 5793.00 | 180.16 | N/A |
| 1-Kestose | oligosaccharide (positive control) | KES | 440080.00 | 504.40 | N/A |
| Fructooligosaccharide | oligosaccharide (positive control) | FOS | N/A | N/A | N/A |

Supplementary Table S1 (Continued)

| Sweetener* | Supplier and catalog number | Acceptable Daily Intake (ADI) (mg/kg bw/d) | cADI without considering proportion reaching the colon (mg/mL) | Proportion that reaches the colon (%) | Concentration in medium (mg/mL) |
| --- | --- | --- | --- | --- | --- |
| Acesulfame K | TCI A1490 | 15 (FDA) | 0.05 | 1^1^ | 0.05 (AE005) and 2 (ACE2) |
| Aspartame | Alfa Aesar J61523 | 50 (FDA) | 17.58 | 0^2^ | 2.00 |
| Advantame | Sigma-aldrich 80054 | 32.8 (FDA) | 10.31 | 89.5^3^ | 2.00 |
| Neotame | Sigma-aldrich 49777 | 0.3 (FDA) | 0.07 | 63.7^4^ | 0.067 (NEO006) and 2 (NEO2) |
| Saccharin (Sodium saccharin dihydrate) | J&K 926097 | 15 (FDA) | 0.53 | 10^5^ | 0.5 (SAC05) and 2 (SAC2) |
| Sucralose | Alfa Aesar J66736 | 5 (FDA) | 1.49 | 85^6^ | 1.50 |
| Stevia extract (Stevioside) | TCI S0594 | 4 (FDA) | 1.41 | 100^5^ | 1.40 |
| Stevia extract (Rebaudioside A) | TCI R0095 | 4 (FDA) | 1.41 | 100^5^ | 1.40 |
| Monk fruit extract | Sigma USP 1445492 | 6.8 (FDA)** | 2.39 | 100^5^ | 2.40 |
| Thaumatin | TCI T1144 | 1.1 (FDA)*** | 0.39 | 0^7^ | 0.39 (THA03) and 2 (THA2) |
| Cyclamate (Sodium cyclamate) | Alfa Aesar A18666 | 7 (EFSA) | 1.55 | 63^8^ | 1.60 |
| Neohesperidin Dihydrochalcone | TCI N0675 | 5 (EFSA) | 1.76 | 100^9^ | 1.80 |
| Salt of Aspartame-Acesulfame | Sigma USP 1043750 | 20.46 (EFSA)**** | 7.19 | 0^8^ | 2.00 |
| Sorbitol (D-Sorbitol) | TCI S0065 | N/D | N/A | 75^10^ | 2.00 |
| Mannitol (D-Mannitol) | TCI M0044 | N/D | N/A | 75^10^ | 2.00 |
| Isomalt | Sigma-aldrich PHR1769 | N/D | N/A | 90^10^ | 2.00 |
| Maltitol | TCI M0797 | N/D | N/A | 60^10^ | 2.00 |
| Lactitol (Lactitol monohydrate) | J&K 126721 | N/D | N/A | 98^10^ | 2.00 |
| Xylitol | TCI X0018 | N/D | N/A | 50^10^ | 2.00 |
| Erythritol (meso-Erythritol) | TCI E0021 | N/D | N/A | 10^10^ | 2.00 |
| Hydrogenated starch hydrolysates | CarboMer Inc. 68425-17-2 | N/D | N/A | 60^10^ | 2.00 |
| Glucose (D-Glucose) |  | N/A | N/A | N/A | 2.00 |
| 1-Kestose |  | N/A | N/A | N/A | 2.00 |
| Fructooligosaccharide |  | N/A | N/A | N/A | 2.00 |

*Names of the compounds that were used to represent the sweetener are shown in the brackets.

**The estimated 90th percentile intake of MFE in the general population by the FDA is used here, as its ADI is not specified^11^.

*** The highest estimated exposure level of THA in the general population by the EFSA is used here, as its ADI is not specified^12^.

****The ADI of SAA is calculated based on ADI of both ASP and acesulfame, as specified by the EFSA^13^.

**Supplementary Table S2** Functional annotation of enriched discriminative proteins from Clostridia.

| Protein Group | COG Category | Top1 Protein ID | COG | Functions |
| --- | --- | --- | --- | --- |
| CHO elevated | C | MH0088_GL0108545 | COG0427 | Acyl-CoA hydrolase |
| CHO elevated | C | MH0170_GL0056715 | COG1145 | Ferredoxin |
| CHO elevated | C | MH0012_GL0011876 | COG1951 | Tartrate dehydratase alpha subunit/Fumarate hydratase class I, N-terminal domain |
| CHO elevated | C | MH0062_GL0059388 | COG0055 | FoF1-type ATP synthase, beta subunit |
| CHO elevated | C | MH0006_GL0217568 | COG1145 | Ferredoxin |
| CHO elevated | C | MH0020_GL0049440 | COG4656 | Na+-translocating ferredoxin:NAD+ oxidoreductase RNF, RnfC subunit |
| CHO elevated | C | MH0086_GL0053161 | COG1148 | Heterodisulfide reductase, subunit A (polyferredoxin) |
| CHO elevated | C | MH0086_GL0053163 | COG2048 | Heterodisulfide reductase, subunit B |
| CHO elevated | C | MH0086_GL0064754 | COG0282 | Acetate kinase |
| CHO elevated | C | MH0087_GL0025781 | COG0055 | FoF1-type ATP synthase, beta subunit |
| CHO elevated | C | MH0087_GL0028786 | COG0479 | Succinate dehydrogenase/fumarate reductase, Fe-S protein subunit |
| CHO elevated | C | MH0131_GL0153314 | COG1053 | Succinate dehydrogenase/fumarate reductase, flavoprotein subunit |
| CHO elevated | C | MH0131_GL0154648 | COG1838 | Tartrate dehydratase beta subunit/Fumarate hydratase class I, C-terminal domain |
| CHO elevated | C | MH0131_GL0154649 | COG1951 | Tartrate dehydratase alpha subunit/Fumarate hydratase class I, N-terminal domain |
| CHO elevated | C | MH0200_GL0037813 | COG1145 | Ferredoxin |
| CHO elevated | C | MH0205_GL0101334 | COG1145 | Ferredoxin |
| CHO elevated | C | MH0252_GL0259156 | COG1145 | Ferredoxin |
| CHO elevated | C | SZEY-41A_GL0041857 | COG0056 | FoF1-type ATP synthase, alpha subunit |
| CHO elevated | C | V1.FI36_GL0193889 | COG1908 | Coenzyme F420-reducing hydrogenase, delta subunit |
| CHO elevated | J | MH0122_GL0093568 | COG0244 | Ribosomal protein L10 |
| CHO elevated | J | MH0147_GL0131451 | COG0539 | Ribosomal protein S1 |
| CHO elevated | J | 742740.HMPREF9474_00428 | COG0094 | Ribosomal protein L5 |
| CHO elevated | J | MH0004_GL0001738 | COG0186 | Ribosomal protein S17 |
| CHO elevated | J | MH0006_GL0034105 | COG0203 | Ribosomal protein L17 |
| CHO elevated | J | MH0074_GL0039008 | COG1841 | Ribosomal protein L30/L7E |
| CHO elevated | J | MH0092_GL0132892 | COG0256 | Ribosomal protein L18 |
| CHO elevated | J | MH0161_GL0124112 | COG0200 | Ribosomal protein L15 |
| CHO elevated | J | MH0188_GL0186972 | COG0480 | Translation elongation factor EF-G, a GTPase |

| Protein Group | COG Category | Top1 Protein ID | COG | Functions |
| --- | --- | --- | --- | --- |
| CHO elevated | J | MH0196_GL0121197 | COG0103 | Ribosomal protein S9 |
| CHO elevated | J | O2.CD1-0-PT_GL0056648 | COG0441 | Threonyl-tRNA synthetase |
| CHO elevated | J | T2D-19A_GL0015282 | COG1544 | Ribosome-associated translation inhibitor RaiA |
| CHO elevated | O | MH0198_GL0077222 | COG0544 | FKBP-type peptidyl-prolyl cis-trans isomerase (trigger factor) |
| CHO elevated | O | BGI003A_GL0018079 | COG0234 | Co-chaperonin GroES (HSP10) |
| CHO elevated | O | MH0020_GL0059731 | COG0576 | Molecular chaperone GrpE (heat shock protein) |
| CHO elevated | O | MH0086_GL0106459 | COG0326 | Molecular chaperone, HSP90 family |
| CHO elevated | O | MH0212_GL0048164 | COG0822 | NifU homolog involved in Fe-S cluster formation |
| CHO elevated | O | MH0259_GL0189842 | COG0652 | Peptidyl-prolyl cis-trans isomerase (rotamase) - cyclophilin family |
| CHO depleted | C | 457421.CBFG_01667 | COG3411 | (2Fe-2S) ferredoxin |
| CHO depleted | C | 457421.CBFG_03929 | COG1251 | NAD(P)H-nitrite reductase, large subunit |
| CHO depleted | C | MH0205_GL0028253 | COG1882 | Pyruvate-formate lyase |
| CHO depleted | C | MH0100_GL0117974 | COG1454 | Alcohol dehydrogenase, class IV |
| CHO depleted | C | MH0100_GL0136349 | COG5016 | Pyruvate/oxaloacetate carboxyltransferase |
| CHO depleted | C | MH0116_GL0144625 | COG0538 | Isocitrate dehydrogenase |
| CHO depleted | C | MH0131_GL0135388 | COG0554 | Glycerol kinase |
| CHO depleted | C | MH0131_GL0144928 | COG1251 | NAD(P)H-nitrite reductase, large subunit |
| CHO depleted | C | MH0193_GL0111985 | COG1454 | Alcohol dehydrogenase, class IV |
| CHO depleted | C | MH0086_GL0013793 | COG1454 | Alcohol dehydrogenase, class IV |
| CHO depleted | C | BGI-06A_GL0041136 | COG1145 | Ferredoxin |
| CHO depleted | C | HT25A_GL0108527 | COG0554 | Glycerol kinase |
| CHO depleted | C | MH0108_GL0033324 | COG1156 | Archaeal/vacuolar-type H+-ATPase subunit B/Vma2 |
| CHO depleted | C | MH0148_GL0067799 | COG5016 | Pyruvate/oxaloacetate carboxyltransferase |
| CHO depleted | C | MH0230_GL0024017 | COG1052 | Lactate dehydrogenase or related 2-hydroxyacid dehydrogenase |
| CHO depleted | C | MH0239_GL0010131 | COG1156 | Archaeal/vacuolar-type H+-ATPase subunit B/Vma2 |
| CHO depleted | C | MH0239_GL0021769 | COG1251 | NAD(P)H-nitrite reductase, large subunit |
| CHO depleted | C | V1.CD12-3_GL0086554 | COG4577 | Carboxysome shell and ethanolamine utilization microcompartment protein CcmL/EutN |
| CHO depleted | E | 457421.CBFG_04540 | COG0834 | ABC-type amino acid transport/signal transduction system, periplasmic component/domain |

**Supplementary Table S2** (*Continued*)

**Supplementary Table S2** (*Continued*)

| Protein Group | COG Category | Top1 Protein ID | COG | Functions |
| --- | --- | --- | --- | --- |
| CHO depleted | E | 457421.CBFG_05366 | COG0683 | ABC-type branched-chain amino acid transport system, periplasmic component |
| CHO depleted | E | HT25A_GL0006284 | COG0834 | ABC-type amino acid transport/signal transduction system, periplasmic component/domain |
| CHO depleted | E | MH0020_GL0057779 | COG0624 | Acetylornithine deacetylase/Succinyl-diaminopimelate desuccinylase or related deacylase |
| CHO depleted | E | MH0100_GL0157370 | COG0137 | Argininosuccinate synthase |
| CHO depleted | E | MH0131_GL0038390 | COG0747 | ABC-type transport system, periplasmic component |
| CHO depleted | E | MH0131_GL0158083 | COG0136 | Aspartate-semialdehyde dehydrogenase |
| CHO depleted | E | MH0227_GL0044967 | COG0059 | Ketol-acid reductoisomerase |
| CHO depleted | E | MH0270_GL0105277 | COG0334 | Glutamate dehydrogenase/leucine dehydrogenase |
| CHO depleted | E | MH0358_GL0146559 | COG0334 | Glutamate dehydrogenase/leucine dehydrogenase |
| CHO depleted | E | T2D-61A_GL0014321 | COG0112 | Glycine/serine hydroxymethyltransferase |
| CHO depleted | E | T2D-61A_GL0044107 | COG0834 | ABC-type amino acid transport/signal transduction system, periplasmic component/domain |
| CHO depleted | G | 457421.CBFG_04444 | COG1653 | ABC-type glycerol-3-phosphate transport system, periplasmic component |
| CHO depleted | G | T2D-61A_GL0015555 | COG2376 | Dihydroxyacetone kinase |
| CHO depleted | G | 457421.CBFG_03875 | COG0057 | Glyceraldehyde-3-phosphate dehydrogenase/erythrose-4-phosphate dehydrogenase |
| CHO depleted | G | MH0131_GL0042674 | COG0696 | Phosphoglycerate mutase (BPG-independent, AlkP superfamily) |
| CHO depleted | G | MH0187_GL0022290 | COG1264 | Phosphotransferase system IIB components |
| CHO depleted | G | N034A_GL0057906 | COG0297 | Glycogen synthase |
| CHO depleted | G | MH0089_GL0008731 | COG1455 | Phosphotransferase system cellobiose-specific component IIC |
| CHO depleted | G | MH0141_GL0047226 | COG1080 | Phosphoenolpyruvate-protein kinase (PTS system EI component in bacteria) |
| CHO depleted | G | MH0211_GL0047626 | COG5263 | Glucan-binding domain (YG repeat) |
| CHO depleted | G | MH0002_GL0043706 | COG0191 | Fructose/tagatose bisphosphate aldolase |
| CHO depleted | G | MH0087_GL0016064 | COG0126 | 3-phosphoglycerate kinase |
| CHO depleted | G | MH0389_GL0105436 | COG0580 | Glycerol uptake facilitator and related aquaporins (Major Intrinsic Protein Family) |

(2)	B, C.; Trugo, L.; P, F. Encyclopedia of Food Sciences and Nutrition.; 2003.

(8)	L., O.-N. Alternative Sweeteners.

(9)	SCIENTIFIC OPINION Flavouring Group Evaluation 32 (FGE.32): Flavonoids (Flavanones and Dihydrochalcones) from Chemical Groups 25 and 30. EFSA Journal No. 2010; 8(9):1065. https://doi.org/10.2903/j.efsa.2010.1065.

(10)	Livesey, G. Health Potential of Polyols as Sugar Replacers, with Emphasis on Low Glycaemic Properties. Nutr. Res. Rev. 2003, 16 (2), 163–191. https://doi.org/10.1079/NRR200371.

(11)	GRAS Notice 627: Siraitia Grosvenorii Swingle (Luo Han Guo) Fruit Juice Concentrate; 2016.

(12)	Canfora, E. E.; Jocken, J. W.; Blaak, E. E. Short-Chain Fatty Acids in Control of Body Weight and Insulin Sensitivity. Nat Rev Endocrinol 2015, 11 (10), 577–591. https://doi.org/10.1038/nrendo.2015.128.

(13)	Hamer, H. M.; Jonkers, D.; Venema, K.; Vanhoutvin, S.; Troost, F. J.; Brummer, R.-J. Review Article: The Role of Butyrate on Colonic Function: REVIEW: ROLE OF BUTYRATE ON COLONIC FUNCTION. Alimentary Pharmacology & Therapeutics 2007, 27 (2), 104–119. https://doi.org/10.1111/j.1365-2036.2007.03562.x.
